# Mixed evidence for intralocus sexual conflict from male-limited selection in *Drosophila melanogaster*

**DOI:** 10.1101/2024.10.12.618000

**Authors:** Harshavardhan Thyagarajan, Imran Sayyed, Mindy G. Baroody, Joshua A. Kowal, Troy Day, Adam K. Chippindale

## Abstract

Sexual conflict over shared traits – intralocus sexual conflict (IaSC) – may be common and consequential, but experimental tests of its relative magnitude are challenging and limited in number. We use a sex-limited selection experiment, designed to subject genomic haplotypes of *Drosophila melanogaster* to selection for male fitness without opposing selection acting on female fitness. Importantly, we use three novel base populations to compare results with those from the LH_M_ population, the sole population investigated using this technique. In contrast with previous studies, we find that male fitness of haplotypes subject to male-limited selection (ML populations) are not consistently better than their matched (MC) controls when tested in the “wildtype” state. Males from ML lines did not outperform controls in competitive fitness assays, mate choice trials, fecundity induction or sperm offense tests. As predicted, genetic variation for male fitness was reduced, with low fitness haplotypes apparently removed by selection, but this was only surveyed in one replicate population pair and included a potential artefact in the protocol. Female fitness was markedly reduced by carriage of ML genomic haplotypes, as predicted by sexual antagonism. Hence, our results are only partially consistent with the IaSC hypothesis, raising questions about the relative contribution of sexual conflict to the standing genetic variation in these populations and the potential role of artefacts in the protocol that may have obscured our ability to detect IaSC.

## INTRODUCTION

The evolution of anisogamy is hypothesized to have been a key driver in the origin of sex-specific selection pressures, resulting in different morphological, life-history, reproductive and ecological strategies in a wide range of taxa. To achieve sex-specific adaptation despite a shared genome, a number of genomic mechanisms, such as gene duplications, sex chromosomes, imprinting, and in cis-regulation have evolved in concert with hormones and other signaling systems. Despite these means to facilitate disruptive selection, many sexually-dimorphic traits show positive genetic correlations between the sexes (Poissant *et al*. 2010). These correlations are expected to act as constraints upon the evolution of further dimorphic responses to selection. This brake on the ability to evolve to separate fitness optima for shared traits results in a conflict between the sexes, often labelled *intralocus sexual conflict* (IaSC). Anecdotal evidence for IaSC includes the expression of ornaments and armaments in the sex that does not deploy them in sexual selection (e.g., horns in female soay sheep) suggesting the evolution of dimorphism is sometimes incomplete.

There is a longstanding theoretical interest in IaSC and the conditions under which polymorphism for sexually antagonistic alleles can be sustained by selection (e.g., Kidwell *et al*. 1977; Parker 1979, Lande 1980). For example, Kidwell *et al*. (1977) showed that the degree of dominance and its sex-specificity plays a key role in determining equilibrium behaviour for alleles with opposing fitness effects upon the sexes. Subsequently Rice (1984) recognized the potential role of sex chromosomes, which present distinctive features in XY and ZW systems: specifically, sex-limitation of the Y/W and haplodiploidy of the X/Z chromosomes. The sex-determining chromosomes (Y/W) have been theorized to attract male/female benefit SA loci and make them sex-limited in the non-recombining region, a central hypothesis for the origin and evolutionary dynamics of these chromosomes (Charlesworth & Charlesworth 2000). Assuming identical allelic dominance parameters in males and females, Rice (1984) predicted an enrichment of SA variation on the haplo-diploid X/Z chromosome. The logic of Rice’s model is that dominant female-benefit alleles can accumulate to appreciable frequencies because of the two-fold advantage females have in X-expression in populations. On the other hand, recessive male-benefit alleles can reach high frequencies because their deleterious effects upon females are masked in the heterozygous state. However contrasting theoretical predictions about the SA loci enrichment on the haplodiploid sex chromosome result from differing assumptions about allele dominances (Fry 2010).

Unlike the theory, the empirical investigation of IaSC is a burgeoning area of investigation. Early experimental evidence for the existence of IaSC was provided by Chippindale *et al*. (2001) in the form of a negative genetic correlation for reproductive fitness between the two sexes in *Drosophila*. This measure of intersexual (or cross-sex) genetic correlation for fitness (r*_w,g,m-f_*) has since been used to investigate the existence of IaSC, with negative measures of r*_w,g,m-f_* treated as evidence (cf. Connallon & Matthews 2019). The intersexual genetic correlation for fitness (or components thereof) has been explored using sex-specific parent-offspring data in natural and semi-natural populations, as well as in controlled breeding designs in the lab (reviewed in Bonduriansky & Chenoweth 2009; Poissant *et al*. 2010; Connallon & Matthews 2019). Besides measurements of intersexual genetic correlations for fitness at the population level, a second key focus of this literature has been on specific traits that are potential fulcrums of sexual antagonism. Through measurements of extant trait values and fitness optima (estimated using selection gradients), a number of dimorphic traits such as body size and development time (amongst others) have been shown to have SA fitness effects. Numerous taxa appear to exhibit IaSC, including species of fruit flies, seed beetles, birds, ground crickets, meal moths, plants, lizards, snakes, voles and red deer (reviewed in Bonduriansky & Chenoweth 2009; Cox & Calsbeek 2009; Singh & Punzalan 2018).

Empirical research has so far suggested that the *Drosophila melanogaster* X chromosome does indeed harbour more SA loci than randomly expected (Gibson *et al*. 2002; Pischedda & Chippindale 2006; Long *et al*. 2012; Ruzicka *et al*. 2019, Wong & Holman 2023, but see Lund-Hansen *et al*. 2020, Abbott *et al*. 2020). However, Ruzicka & Connallon (2020) suggest that quantitative genetic attempts at identifying SA loci are likely to overestimate the enrichment of SA loci on the X chromosome, due to asymmetric inheritance pattern of X chromosomes.

Further, recently unearthed evidence for sex-specific dominance reversals (Grieshop & Arnqvist 2018, Pearse *et al*. 2019, Geeta Arun *et al*. 2021, reviewed in Grieshop *et al*. 2024) contributes weight to Kidwell *et al*.’s assumptions, and predictions of random distribution of SA loci across autosomes and sex chromosomes.

As a form of balancing selection acting upon the genome constitutively, IaSC is expected to maintain polymorphisms and contribute to genetic variance for traits related to fitness (Prout 2000). However, while the conditions for sexually antagonistic selection are believed widespread, the conditions for the maintenance of genetic variation may or may not be met in nature. Rapid changes in the environment can cause maladaptation, which can align the selection pressures acting on the two sexes, but gradual environmental fluctuations are likely to stabilise antagonistic variation (Connallon & Hall 2016). While early models typically assume single-locus traits, Flintham *et al*. (2023) further complicate this picture by explicitly considering continuous traits governed by multiple loci. Here, they find that such variation due to sexual antagonism should arise only rarely and often be transient. There is little empirical work directly supporting the maintenance of genetic variance through SA polymorphisms. Ruzicka *et al*. (2019) show that the SA loci they identify in a lab population display signatures of balancing selection (measured using Tajima’s D and minor allele frequency measurements) and identify similar signatures in distantly related populations of fruit flies, suggesting stable balancing selection over long periods of evolutionary time at these loci. More recently, Kaufmann *et al*. (2023) show that artificial sexually antagonistic selection on body size maintains greater genetic variance than sex-limited selection.

In this study, we look to a sex-limited selection protocol [developed by Rice (1996) and pioneered as a tool to study IaSC by Prasad *et al*. (2007)] to address these questions. Using the lack of molecular recombination in *D. melanogaster* males and an independent population of “clone-generator” (CG) females carrying compound-X chromosomes (X-XY karyotype) and autosomal translocations, we are able to reliably passage haplotypes containing an X chromosome and all major autosomes as “hemiclones” from father to son. Starting with outbred stocks and removed from the influence of selection acting upon the female function, we predicted that these populations of hemiclones would experience a knock-down of IaSC, increasing the frequency of male benefit alleles at polymorphic SA loci.

This male-limited (ML) system holds potential advantages over other contemporary systems used in the literature. While pedigree designs allow us to study the standing genetic variance and intersexual correlations for fitness in a population, they tend to require large numbers of sampled lines. To probe the system past fitness and identify SA polymorphisms on the genome, there is a need for complex statistical manipulations, such as the extraction of an “antagonism index” (Innocenti & Morrow 2010; Hill *et al*. 2017; Ruzicka *et al*. 2019). Further, it is not amenable to examination of other phenotypic trait variances and corresponding analyses.

In contrast, the male-limited evolution approach (1) provides a direct experimental manipulation rather than a correlational analysis, (2) allows for statistical simplicity and strength at smaller sample sizes through the amalgamation of male-benefit alleles across many loci in the same individuals, increasing signal-noise ratio and (3) allows us to probe adaptive responses in the phenotype when released from positive trait intersex genetic correlations.

An unfortunate consequence of a tight knit research community is that most experimental studies of sexual conflict in *Drosophila* have been conducted with a single stock population: the LH_M_ (LH-moderate density) population collected in central California in 1988 by L. Harshman (Rice 1996, Rice 1998, Chippindale *et al*. 2001, Gibson *et al*. 2002, Byrne & Rice 2005, Lew & Rice 2005, Lew *et al*. 2005, Linder & Rice 2005, Pischedda & Chippindale 2006; Prasad *et al*. 2007, Long *et al*. 2007, Bedhomme *et al*. 2008, Rode & Morrow 2009, Innocenti & Morrow 2010, Abbott *et al*. 2013, Jiang et al. 2011, Collet *et al*. 2017, Hill *et al*. 2017, Ruzicka *et al*. 2019, Abbott *et al*. 2020, Lund-Hansen *et al*. 2020, Manat *et al*. 2021, Lund-Hansen *et al*. 2022). While results from Ruzicka *et al*. (2019) suggest some generalizability from LH_M_ to the species at large, it is important to affirm this notion by conducting experimental work on different populations. Interestingly, even with LH_M_, Jiang et al. (2011) found markedly different results from Rice (1996) for male postcopulatory sexual selection, and Collet *et al*. (2017) demonstrate that even recently separated populations (∼ 200 generations) of LH_M_ display differences in sexual antagonism. These disparities may highlight sensitivity of the standing genetic variation to the conditions or duration of maintenance in the laboratory of the very same population, and call for the investigation of other populations to establish generalities about the nature and level of sexual conflict within the species.

We conducted our male-limited selection experiment on three new populations from a different geographic origin (Massachusetts, USA), separated from one another since 1980. We report on the competitive reproductive fitness (CRF) of males and females from selected and control lines. Using an improved breeding design in which we strip test individuals of all artefacts deriving from the selection experiment to assess CRF, and we further examine test males for mating success, fecundity induction, sex ratio drive and sperm offense. Lastly, we use a hemiclonal analysis to study genetic variance for reproductive fitness in males and females from selected and control populations, and use this design to report on the intersex genetic correlation for fitness.

We predicted (1) increased male fitness in selected populations, at the cost of female fitness, (2) masculinization of partially dimorphic traits in both males and females (as seen in Prasad *et al*. 2007, Abbott *et al*. 2013), and (3) a reduction of genetic variance for fitness, particularly in males. Previous work suggested that these ML males would be more attractive (Bedhomme *et al*. 2008), but conflicting results exist for post-copulatory sexual selection.

Because sperm and seminal fluid characteristics are presumed to be already male limited traits (Friberg *et al*. 2005, Bjork *et al*. 2007, Jiang *et al*. 2011), we anticipate that the improvement of male fitness will not evolve through improved fecundity induction or sperm competitive ability (although see Rice 1996, 1998). Previous work reports no precedent of driving X chromosomes, but the selection experiment should strongly select for even weakly driving ML-X chromosomes, should they exist. We also test ML-evolved lines with and without the evolved X chromosome to test the prior claims of it being a “hotspot” for sexually antagonistic loci.

## METHODS

### ML Selection

#### Source populations

Our experiments employed selected and control populations originating from the IV base population derived from the wild in 1975 in South Amherst (MA, USA; see Rose 1984). The source populations have had a somewhat varied history of selection, first as lines selected for postponed senescence, and then as controls for a stress resistance selection experiment since 1989 on a 4-week discrete generation cycle (∼500 generations) before our deployment in the current experiments. Three replicate CO populations (CO_1_, CO_3,_ CO_5_) were arbitrarily chosen from the five replicate CO’s and were maintained on a 2-week cycle (day 1-11 in vials, followed by days 12-14 in ∼ 3 litre population cages with food in a petri-dish) for 24 generations. After 24 generations, a male-limited (ML) line and matched-control (MC) line were derived from each of the three replicate populations. These were correspondingly labelled ML_1_-MC_1_, ML_3_-MC_3,_ and ML_5_-MC_5_. All populations were maintained under standard laboratory conditions: moderate density (larval density of 80 - 100 larvae / vial), 25°C, 12:12 light:dark cycle on banana/agar/killed-yeast medium, and with a census population size of ∼1,000 haploid genomes per generation.

### Selection treatment

ML selection lines were established nearly identically to the protocol described in Prasad *et al*. (2007) (fig. S1a). Briefly, 1,000 males from the replicate base population were mated to “clone-generator” (CG) females that carried a compound X(C(1)DX*, y, f*), a Y chromosome and a homozygous-viable translocation of the two major autosomes (T(2 : 3)*rdgc st in ri p^p^*). Using the genetic markers for eye colour, the absence of molecular recombination in males, and the aneuploid mortality experienced by animals inheriting uneven combinations of translocated autosomes, we select sons carrying a haploid genome (“hemiclone”) from their sire, and repeatedly crossed these sons to CG females to passage these hemiclones under conditions of ML selection. 1,000 such males were collected under light CO_2_ anaesthesia each generation on day 10-11 post oviposition (PO) and were combined on day 12 with virgin CG females of matched age in 3 cages (333-334 males and females in each 2L plastic cage). The introduction of a cage phase is a departure from previous ML-evolution protocols, potentially increasing environmental complexity. Eggs were collected from these cages on day 14 at a density of 320- 400 eggs per vial, to achieve a larval density of 80-100 per vial after accounting for aneuploid viability costs. To match these conditions, 500 males (1000 haploid genomes) from each MC line collected under light CO_2_ anaesthesia each generation on day 10-11 PO and combined on day 12 with virgin MC females in 2 cages (250 pairs per cage). Eggs were collected from these lines at a density of 80-100 per vial, with 20 vials dedicated to females and virgin collection and 20 to male collections.

### Recombination Box

In the absence of molecular recombination in males, and the artificial prevention of independent assortment of chromosomes, the ML selection regime is not only sex-limited, but also asexual in propagation, exposing it to higher levels of genetic hitchhiking. We followed the system described in Prasad *et al*. (2007) and cycled a subset of the ML hemiclones through a two generation cross, where ML-selected chromosomes were concentrated in the F_2_ generation in the female sex as wildtype animals, reintroducing recombined ML haplotypes into the selected populations.

Specifically, each generation we collected 50 ML males from each replicate population and crossed them to virgin CG females carrying an additional dominant eye colour marker (*bw^D^*). Sons derived from this first recombination cross (RC1) carrying haploid genomes of interest are brown eyed due to their single copy of *bw^D^*. In the first generation, 50 sons (RC1 males) were crossed to control females, and daughters carrying two ML haploid genomes (identified through absence of *bw^D^*) from this second recombination cross (RC2) were collected as virgins to establish a line that receives ML genetic content through backcrossing with RC1 males. Within 4 generations of backcrossing these females are expected to be ∼95% ML in genome content, increasing incrementally each generation. In each generation after the first generation of the experiment, 70-80 virgin RC2 females were collected and crossed to 50 RC1 males. 50 male offspring from this cross (RC2 males) carrying haploid genomes of interest (rather than translocated autosomes - identified through absence of *bw^D^*) from RC1 fathers and recombined gametes from RC2 mothers are re-introduced into the selected population from which the RC was derived (fig. S1b). This crossing scheme was meant to limit genetic drift but also provided a means by which to derive ML-evolved genomic haplotypes for experimentation, as noted below.

### Experimental animals

We expressed the target (ML and MC) haploid genomes in male and female flies, with an attempt to produce test animals that avoid all sources of artefact from the ML selection regime, including the reversed inheritance of sex chromosomes, the foreign cytoplasm, the maternally inherited Y chromosome, and the translocated autosomes. To do this, we used males and females produced from the second recombination cross in the ML treatment. Through many generations of backcrossing of ML genetic content into a control derived line, we were able to produce RC2 flies carrying ML genetic content without any known associated artefacts. Thus to create the actual test animals, we used a crossing design between RC2 and MC flies.

From a full factorial crossing design, 4 types of male offspring and 3 types of female offspring can be obtained (fig. S2a). From crosses of MC males and females, we derived our control treatment of test animals (male and female). From RC2 males and females, we can derive flies carrying completely ML type genomes, labelled ML_DD_ (ML double dose). Both directions of the hybrid cross produce identical females, carrying an ML haploid genome. These, like males carrying a single complete haploid genome from the ML treatment are labelled ML_SD_ (ML single dose). While crosses with MC sires and RC2 dams produce ML_SD_ males, crosses between RC2 sires and MC dams produce males carrying an X chromosome from MC populations and a single dose of ML autosomes. These males are labelled ML_SD(a)_. Crossing designs used were not fully factorial in all the experiments reported below, but the labelling scheme indicates the crosses carried out as described here. The different test animals studied were denoted using the variable ‘Treatment’ in analysis.

While fitness assays were conducted using test animals as described above, a slightly different design was used to produce test animals for mate choice, sperm competition, body size and development time assays (fig. S2b). Here, test animals were produced using a cross analogous to the second recombination cross used for generating RC2 males to create both ML and MC test flies. First, ML and MC male haplotypes are captured using CG (*bw^D^*) females. Sons from these crosses carrying the haplotypes are crossed to virgin MC or RC2 females to produce test flies. Here too, as before, the treatments are labelled as per the system above (MC, ML_DD_, ML_SD_, ML_SD(a)_). While mostly identical to the design used for fitness assays, the male test animals from this design carry Y chromosomes from clone generator lines that may have been exposed to feminization or mutation accumulation effects. Due to this oversight, it is possible that there are effects in the phenotypes displayed by the test animals in response to the Y chromosome.

To study genetic variance for fitness in these populations, we created hemiclonal lines (as described in Chippindale *et al*. 2001) using single haploid ML-1 and MC-1 genomes sampled through crosses with CG females. Each individual haplotype was amplified entirely in sons using crosses with CG (*bw^D^*) females; and had its X chromosome passaged to a daughter using crosses with a female carrying MC derived autosomes and balancer X chromosomes (fig S3). These sons and daughters respectively were then crossed to one another to produce test animals carrying ML and MC haplotypes, along with control derived autosomes. Females derived were homozygous for the X chromosome, and males carried the Y chromosome from the CG (*bw^D^*), much like the test animals from the mate choice and sperm competition assays.

Across all assays, test animals were maintained at a larval density of 80-100 animals per vial. Virgin flies were collected 6-8 hrs from eclosion using light anaesthesia between days 8-10 post egg laying. Non virgin test animals were collected on days 10-11.

Fitness, mate choice and sperm competition assays required the use of phenotypically marked competitors. These lines allowed us to identify successful mates and measure competitive siring/damming success by distinguishing the progeny phenotype. To this end, we backcross a recessive eye colour markers into control derived lines to create these competitor populations. Two such populations were created, both combining flies across the different replicate populations of the MCs. In the first, we introduced the pink-peach (*p*^p^) eye colour marker, and in the other we introduced the brown (*bw^1^*) eye colour marker. Collectively, we refer to these lines as recessive competitors – “Cr”.

### Assays

#### Competitive Reproductive Fitness (CRF) assays

CRF for was assayed for both males and females. Following the crossing design, test females were collected as virgins, and test males were collected on day 10-11. On day 12, 30 test animals were combined with 30 Cr individuals of the same sex, and 50 Cr of the opposite sex (both carrying the recessive eye colour marker). These animals were held together in mini cages with ad libitum yeast between days 12-14 from egg, to mimic the reproductive conditions of the selection treatment. For each treatment (within each replicate pair of lines), we assessed 6 such competition arenas (labelled contests in analysis), and collected 6 vials of eggs to measure CRF.

Male CRF was assayed thrice, in generations 50, 64 & 70 of the selection experiment. In generations 50 and 64, only ML_SD_ and MC animals were tested. All four were tested in generation 70. For the experiment conducted in the 70th generation, males were collected as virgins. Female CRF was assayed twice, in generations 48 & 50 of the selection experiment. On both occasions, we only tested ML_SD_ and MC animals.

#### Hemiclonal analysis of fitness

Conducted in generation 78, CRF assays for hemiclonal analysis were designed very similarly to the CRF assays described above for males. The competition size was reduced in response to the increase in number of lines being studied. Here, we combined 4 test males with 4 Cr males and 6 Cr females. For female fitness, 4 virgin test animals were combined with 3 Cr males. Rather than competitive reproductive fitness, female productivity was assayed as a proxy for fitness. In both sexes, up to 5 such contests were set up for each line, depending on the availability of usable test animals. Complete hemiclonal data for both sexes was collected from 76 lines out of an initially sampled 80.

#### Mating success assays

We assayed female mate choice (or male mating success) in generation 66, using progeny eye colour. Following the crossing design, test males were collected on day 10-11. On day 12, single test males were combined with a Cr male, and a Cr virgin female. When one of the two possible pairs begin copulation, the excess male was aspirated out by gently removing the foam plug without disturbing the pair in copula. After the mating ended and animals separated, the remaining male was also removed, and the female was left undisturbed to oviposit for 48 hours under *ad-libitum* yeast conditions. The time of introduction (observation start time), amplexus formation (mating start time) and separation (mating end time) were noted, which provides us measures of mating latency and mating duration. The offspring sired in each vial were sexed and enumerated to test for sex ratio drive effects and male fecundity induction, respectively. The eye colour of the offspring in each vial was noted to identify the sire that successfully mated. For each treatment (within each replicate pair of lines), we assessed 50 such competition vials.

#### Sperm offense assays

Sperm offense (or P2) was assayed (in generation 74) by serially mating Cr females first to Cr males and subsequently to target males. Following the crossing design, test males were collected on day 10-11. On day 12, 12 virgin Cr females with *ad-libitum* yeast nutrition were combined in vials with males from the same population. Matings were observed carefully and almost all females were observed to be in amplexus synchronously. Although unlikely, especially as recently mated females tend to be refractory, the group setup did not completely preclude double-mating; however this was consistent across all the treatments. In all vials, at least all but 2 females were directly observed to have been mated. Using a control treatment where no second male was provided, we found that this system resulted in about 97% of females being fertile. An hour after synchronous matings ended, the flies were gently anaesthetized with CO_2_, and males were discarded. On day 13, 10 test males were combined with females. Second matings were observed carefully to ensure that no single female was multiply mated at this stage. To achieve this, we observed all the vials to note when matings began in these vials and continue observing the vials for a period of ∼3 hours after this point to note the number of matings seen, at the end of which the males were discarded to ensure that the first mated female did not return to a receptive state. We then randomly selected 10 females and transferred them to individual test tubes with *ad-libitum* yeast to oviposit for 24 hours, after which they were discarded. Due to the arbitrary cut-off in the second mating period, dictated by the first pair to copulate in each vial, we did not compare the number of matings achieved by target males in each treatment, instead assessing the proportion of offspring sired where a second mating has successfully taken place.

Offspring from each tube were phenotyped by eye colour and enumerated. For each treatment (within each replicate pair of lines), we assessed 30 such competition vials.

### Statistical Analysis

Statistical analysis was conducted in R version 4.2.3 -- "Shortstop Beagle" (R Core Team 2023). Data was visually assessed for residual normality and using Bartlett’s test for residual heterogeneity. Where generalised linear models were used, residual dispersion was tested using the DHARMa package (Hartig 2022).

Linear mixed models were employed to analyse CRF. Treatment and replicate population were used as fixed factors in the analysis to explain the proportion of offspring sired/dammed by the target individuals. To avoid pseudo-replication, contest was included as a random factor.

Where significant differences were identified, we employed planned contrasts between the treatments in a single replicate population pair with a corrected alpha value.

In the hemiclonal analysis of fitness, we analysed genetic variance for fitness in each selection regime, for each sex, using a random effects model that explained the proportion of offspring sired / dammed by the target individuals, using the hemiclonal line as a random factor. The proportion of variance explained by the random factor line was then compared between selection treatments using an F test. Fitness data from the hemiclonal analysis was also analysed using a mixed model, where selection was used as a fixed factor. Here, we did not include replicate as in the case of CRF, as the study was conducted only on replicate population pair 1. Here too, line was included as a random factor. Additionally, mean estimates of male fitness and female fitness for each line from each selection regime were analysed using a Pearson correlation test to test if the ML population displayed a different intersex genetic correlation for fitness from the control. The correlation coefficients were compared against one another using a Fisher’s Z transformation.

To study the mating success of target males in female mate choice trials, we used a generalized linear model with a binomial distribution. The proportion of mating successes for target animals was explained using selection and replicate as fixed factors. Likewise, log(mating latency), mating duration, fecundity induction (offspring count) and sex drive (offspring sex ratio) were analysed using a Gaussian distribution for tubes, with selection and replicate as fixed factors.

For sperm offense, the proportion of offspring sired by the second male was analysed in two steps, as if in a Hurdle model. First, to create a data transformation that enabled the analysis, we subtracted the proportion of offspring sire from 1. This inverted measure was then separated in two parts – values of zero (where all the offspring are sired by the target male) and non-zero values. The number of zeroes attributable to each treatment was analysed as a generalized linear model with a binomial distribution, using treatment and replicate as fixed factors.

## RESULTS

### Competitive reproductive fitness (CRF)

#### Male CRF

In generations 50 and 64, we found no effect of treatment, replicate or their interaction on male CRF for the ML_SD_ treatment, which is most representative of the selection treatment (tableA1a). In generation 50, ML_SD_ displayed a CRF of 0.34 +/- 0.01 (mean +/- 1.96*se) compared to 0.32 +/- 0.01 for MC (fig. 1a). In generation 64, these figures were 0.34 +/- 0.02 and 0.32 +/- 0.02, respectively (fig. 1b).

**Figure 1.**
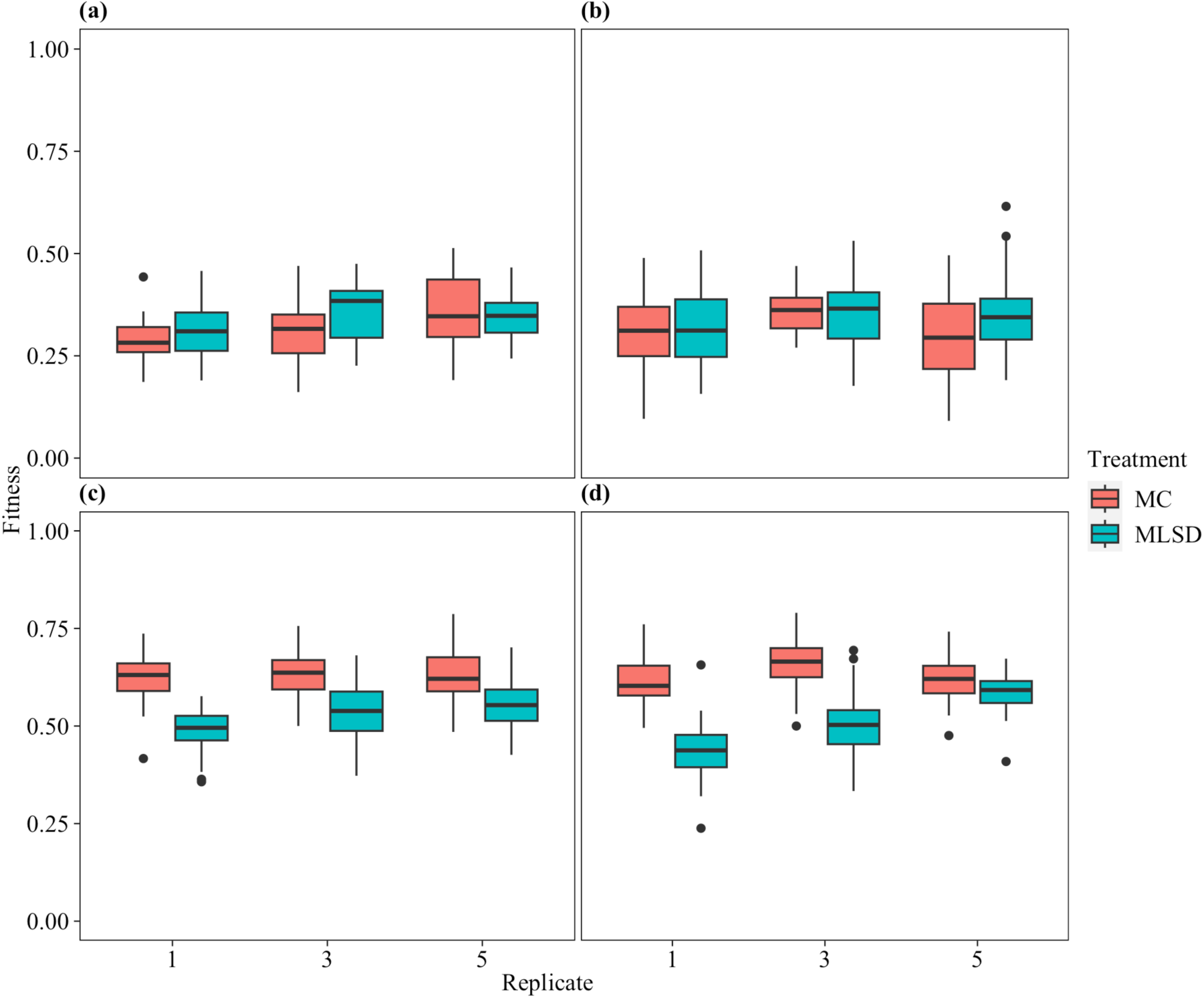
The proportion of offspring (CRF) of target MC (red) & ML_SD_ (blue) males in (a) generation 50, and (b) 64; target females in (c) generation 48, and (d) 50. ML lines were tested as whole (X, II, III) haplotypes in a single copy

In generation 70, we found a significant effect of treatment on reproductive fitness (p < 0.001), and no effect of replicate or their interaction term (although the interaction term approached the α = 0.05 threshold) (table S1b). Excluding replicate (and the corresponding interaction term), we conducted pairwise contrasts between all 4 treatments to find that the only significantly different pairs were ML_SD_ > ML_DD_ (Tukey adjusted p = 0.006) and MC > ML_DD_ (Tukey adjusted p = 0.01). CRF for each treatment (fig 2.2) was as follows: MC (0.45+/-0.02), ML_SD_ (0.47+/-0.02), ML_SD(a)_ (0.42+/-0.02), ML_DD_ (0.37+/-0.02).

**Figure 2.**
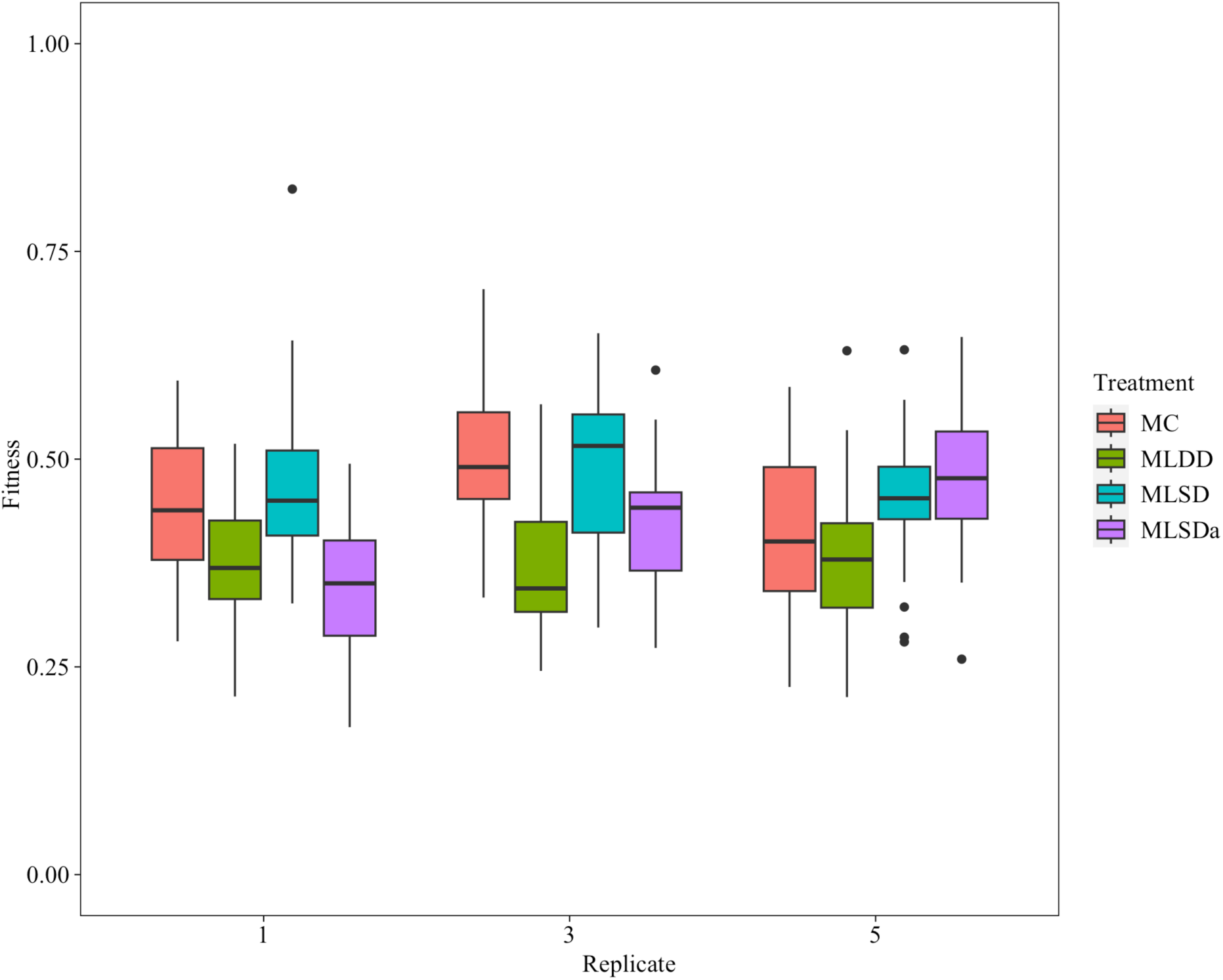
The proportion of offspring sired by target males in generation 70. ML lines were tested as whole (X, II, III) haplotypes in a single copy (ML_SD_ - blue), with just the autosomes in single copy (ML_SD(a)_ - purple) or with all autosomes from the ML populations, a “double-dose” (ML_DD_ - green); see text for details.

#### Female CRF

In generation 48 we found a significant effect of selection treatment (p < 0.0001) on CRF, and no effect of replicate or their interaction term on CRF (table A3a). ML_SD_ females displayed a CRF of 0.53+/-0.01, and MC females 0.63+/-0.01 (fig. 1c).

In generation 50, we found a significant effect of selection treatment (p < 0.0001), replicate (p < 0.0001) and the interaction term (p < 0.001) on CRF (table A3a). Here too, ML_SD_ females displayed reduced CRF compared to MC animals. Contrasts between treatments within each replicate revealed that the difference between ML_SD_ and MC females of replicate 5 was not significant (p = 0.074) at the adjusted α = 0.017 threshold for 3 comparisons. Here too, the ML_SD_ mean was found to be lower than that of MC. Overall, ML females displayed a CRF of 0.51+/- 0.02, and MC females 0.63+/-0.01 (fig. 1d).

### Hemiclonal analysis of fitness

#### Male fitness

We quantified genetic variance for fitness for both ML (0.0509) and MC (0.1771) males as the proportion of variance attributed to ‘line’ in a random effects model. Using an F(37, 37) test, we found that these variances were significantly different from one another (p = 0.0001) (table A2a). Male fitness analysed using selection as a fixed factor and line as a random factor revealed an effect of selection (p < 0.0001) (table A2c). This model did violate the Bartlett’s test for homogeneity of variances, but as shown above, that is likely a feature rather than a bug here. Overall, ML males displayed a CRF of 0.55 +/- 0.03, compared to MC males 0.45 +/- 0.04 (fig 3).

**Figure 3.**
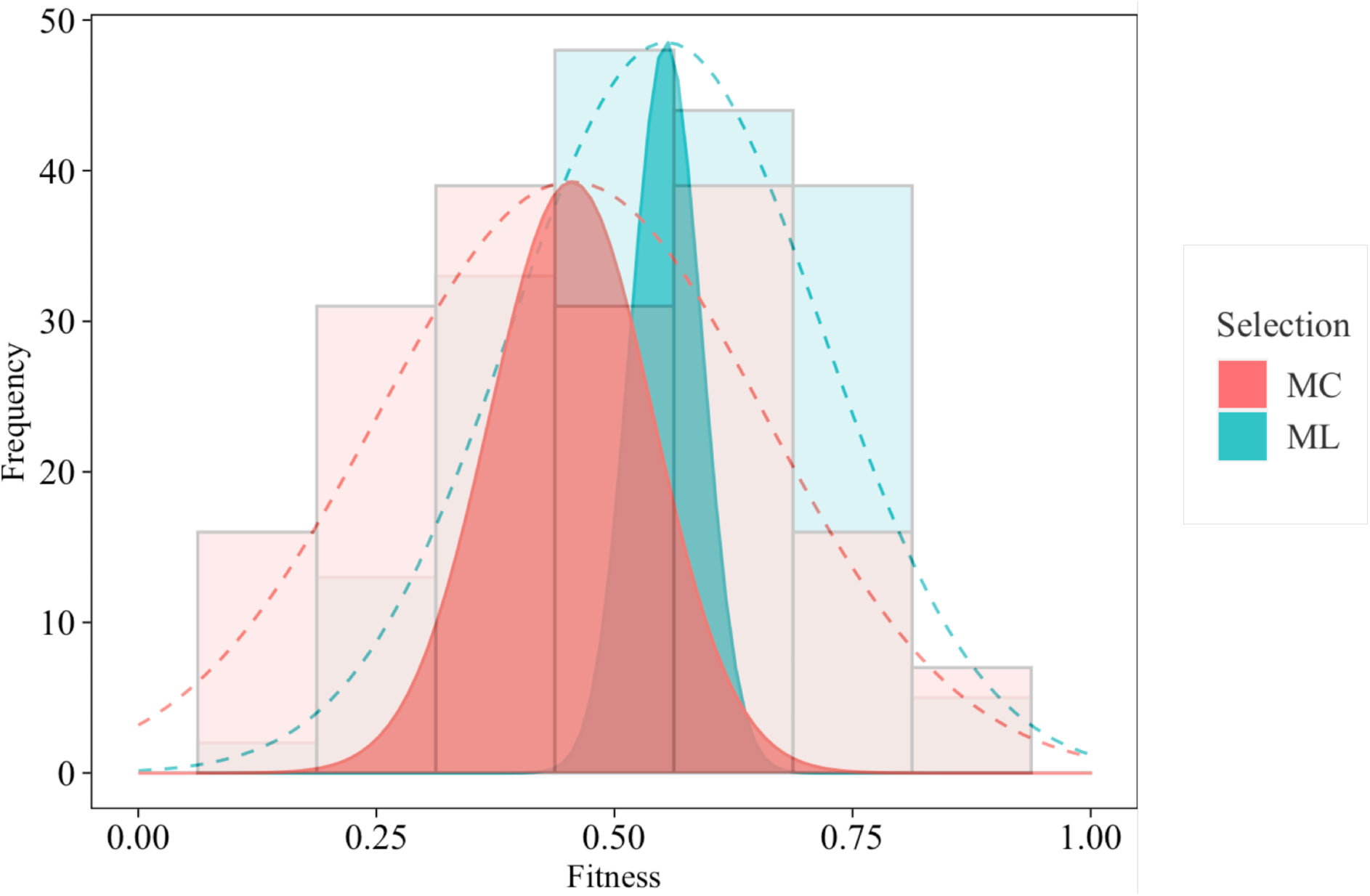
Heritable variance for male CRF. Histograms represent actual measurements from hemiclonal lines from ML (blue) & MC (red) treatments. Mean, variance for each treatment is depicted using dotted curves. Solid distributions depict heritable variance.

#### Female productivity

Likewise, we quantified genetic variance for productivity in ML (0.467) and MC (0.424) females. These were not significantly different from one another in a F(37,37) test (p = 0.39) (table S2b). Female productivity analysed using selection as a fixed factor and line as a random factor revealed no effect of selection (p = 0.29) (table S2c). ML dam productivity was 17.69 +/- 2.05 and for MCs it was 16.09 +/- 1.92.

#### Intersexual genetic correlation for fitness

Intersex genetic correlations for fitness for both the ML (ρ = 0.0247) and MC (ρ = 0.11) population were not significantly different from 0 (fig. 4, table S2d). Likewise, they were not significantly different from one another (p = 0.34) (table S2e).

**Figure 4.**
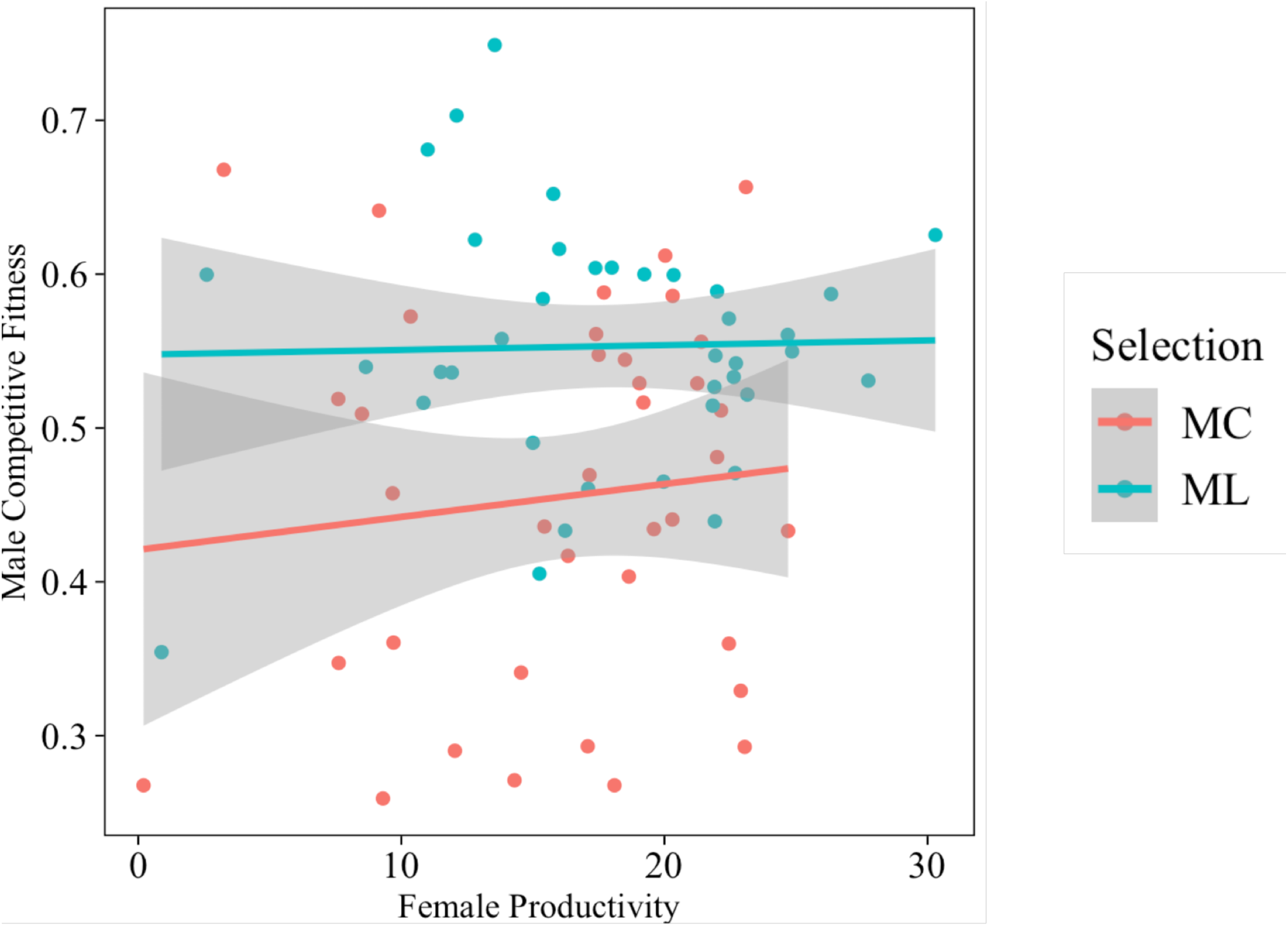
Intersex genetic correlations plotted between female productivity and male CRF for each hemiclonal line from ML (blue) and MC (red) treatments. A linear regression is plotted for each treatment. Shaded region depicts 95% confidence interval.

### Mate choice assays

#### Mate choice

We find no effect of selection, replicate or their interaction upon the male chosen for mating, mating latency or mating duration (tables S4a, b, c). Across all replicates however, we note a consistent direction of difference in mate choice success. ML males were more successful, with greater differences in replicates 1 (61.7% success for ML vs 50.0% MC) and 3 (56.5% vs 46.5%), but also in 5 (44.7% vs 40.0%) (fig 5a). It is possible that there exists an effect the study was not sufficiently powered to capture. MLs on average showed a mating latency of 13.96 +/- 4.14 mins and a mating duration of 18.43 +/- 1.00 mins. For the MCs, these figures were 14.93 +/- 4.33 mins and 17.95 +/- 1.48 mins respectively (fig. 6a, b).

**Figure 5.**
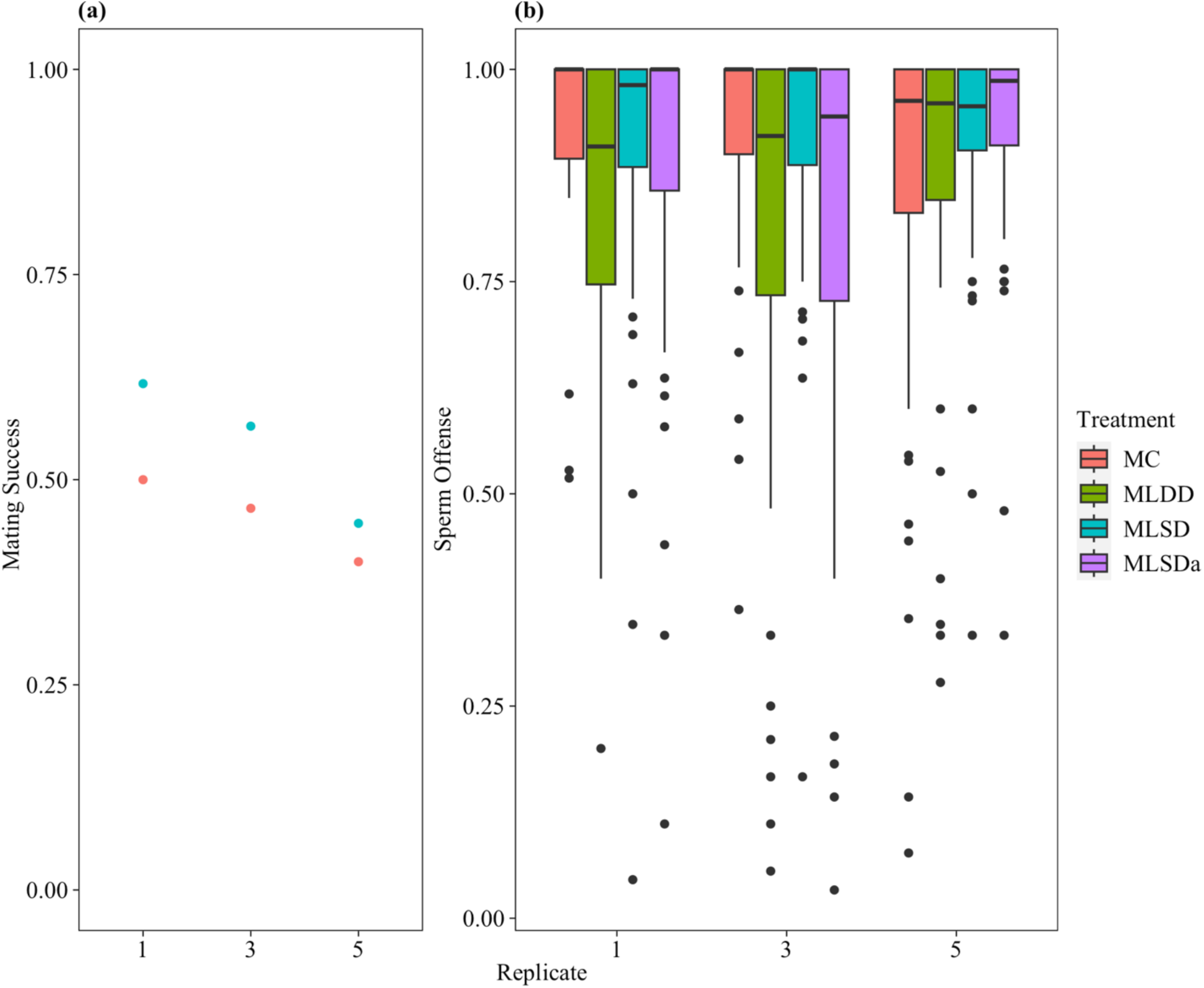
(a) The proportion of target MLSD (blue) or MC (red) males gaining mating with females relative to a marked competitor line and (b) post-copulatory reproductive success as second of two mates (i.e., P2 or sperm offence). ML lines were tested as whole (X, II, III) haplotypes in a single copy (ML_SD_ - blue), with just the autosomes in single copy (ML_SD(a)_ - purple) or with all autosomes from the ML populations, a “double-dose” (ML_DD_ - green); see text for details.

**Figure 6.**
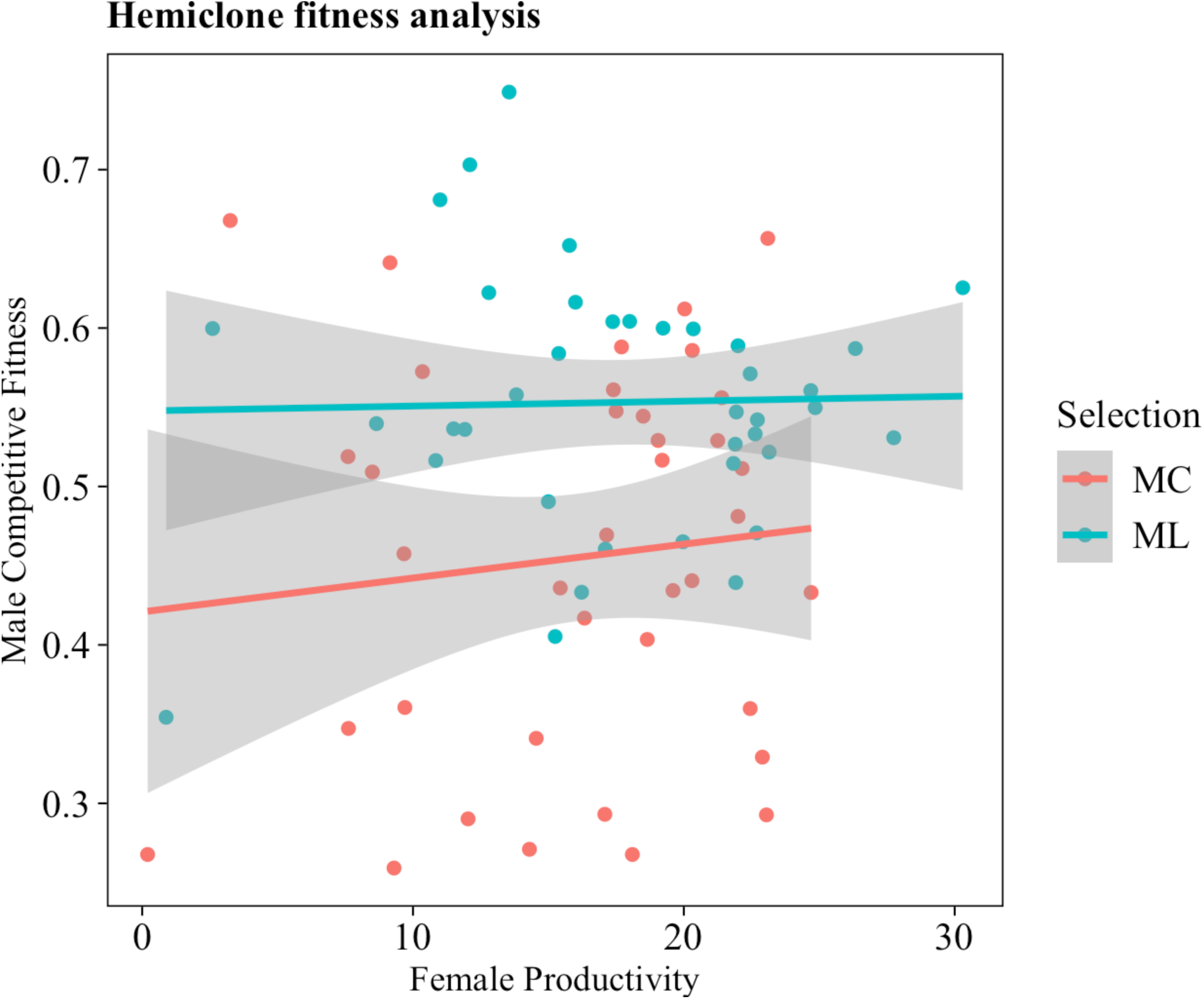
Measures of (a) latency to mating (log transformed), (b) duration of matings, (c) brood sex ratios and (d) fecundity induction for target ML_SD_ (blue)and MC (red) males.

The model analysing mating duration deviated from homogeneity of variances as tested by Bartlett’s test, suggesting a potentially flawed analysis. However, from data visualization it appears that there are no real differences in this dataset, as suggested by the model.

#### Fecundity initiation, sex ratio drivers

We find no effect of selection, replicate or their interaction on fecundity induction, or sex ratio drivers (tables S4d, e). Sex ratio for all the populations hovered around the expected 0.5 mark. ML male offspring were at 0.51 +/- 0.02 and for MCs it was 0.50 +/- 0.03 (fig. 6c). ML males induced a fecundity of 51.09 +/- 2.94 offspring in partners, while MC males achieved 50.94 +/- 3.21 (fig. 6d).

### Sperm offense

We find no effect of selection treatment, replicate or their interaction on either of the sequential steps of analysis on sperm competition (tables 5a, b). Proportion of offspring sired by each treatment as a second male was as follows: MC (0.90+/-0.03), ML_DD_ (0.85+/-0.03), ML_SD(a)_ (0.89+/-0.03), and ML_SD_ (0.90+/-0.02) (fig. 5b).

## DISCUSSION

Sex-limited evolution has been a powerful tool in the study of sexual conflict in *Drosophila melanogaster* but, to our knowledge, has only been used with a single base population – the LH_M_ population originating in central California. Our study, employing populations from a different geographic origin (Massachusetts, USA), fails to find strong evidence for a key proof of principle prediction of this system: the generic improvement of male fitness in response to the male-limited (ML) selection treatment. We find only inconsistent and potentially conditional increases in male competitive reproductive fitness, and no measureable improvements in several of its components. On the other hand, through a hemiclonal analysis of genetic variance in one population pair, we found a significant fitness advantage to the ML-selected population, coupled with a marked reduction in heritable variation for male fitness in the selected population relative to its paired control. Moreover, also consistent with intralocus sexual conflict (IaSC), we found that the fitness of females carrying ML-evolved haplotypes had declined sharply relative to controls, as predicted by selection upon sexually antagonistic variation. Our results compel us to consider that the complex genetic milieu of the selection treatment employed, and the evolutionary history of the populations used in the selection experiment, may have contributed to the mixed findings reported herein.

Previous applications of male-limited selection have documented fitness gains over short evolutionary time-frames, suggesting ample standing genetic variation in the LH_M_ population. Rice (1996) primarily attributed increased male fitness to the adaptation of males to the non-evolving “clone generator” females used to create sex-limited gene transmission. The females, drawn each generation from a separate stock, could not evolve in response to their mates, potentially acted as a “non-responding target”. Hence males that could gain matings with these specific females, or favourably influence their paternity after mating, would be selected. In a subsequent paper (Rice 1998) the potential role of IaSC in the response was recognized, with females expressing ML-evolved hemiclones showing longer development time, which was interpreted as a correlate of reduced fitness. Later work (Prasad et al. 2007) as well as our own recent (*in prep*) results also suggest that development time can evolve in the direction of extant sexual dimorphism under ML-selection, with both sexes slowing. While Prasad et al. (2007) showed evidence for improvement in male fitness from the application of the ML selection regime and unambiguous declines in female fitness, male fitness increases were not tested in the absence of “target” clone-generator females. These results highlight the potential difficulties of discriminating between IaSC and fitness gains through specialization representative of *interlocus* sexual conflict (IeSC).

Our results differ from those earlier outcomes in a number of ways. Most importantly, our competitive reproductive fitness (CRF) assays turned up only small and nonsignificant differences between controls (MC populations) and the ML genetic configuration most representative of the selection environment: X-autosome “hemiclones” paired with MC autosomes (i.e., in the heterozygous state, designated ML_SD_ for “single dose”). Single-dose autosomes in the absence of the ostensible “hotspot” X (ML_SD(a)_) were suggestively lower than ML_SD_ males in fitness in two replicates, but this difference did not stand up to statistical scrutiny. The only difference that did show significance was that ML males carrying a “double dose” of ML autosomes (ML_DD_) performed more poorly than ML males with a single dose (ML_SD(a)_) or

MC controls. This result is intriguing because any ML autosomal adaptation would have to occur in the heterozygous state, and perhaps in response to specific properties of the T2;3 translocation used during selection. Expressed in two copies, ML-evolved chromosomes actually depress male fitness, potentially representing an “overdose” of compensatory factors for deficiencies of the translocation. Related to this, it is also possible that selected autosomes are effectively recessive when paired with wildtype control (MC) autosomes in males.

The lack of measurable differentiation in the male fitness experiments is all the more perplexing given the clarity and consistency of the results for females. In keeping with predictions for IaSC, the CRF of females expressing ML hemiclonal genomes was found to be markedly lower than MC female fitness (minus 15 – 20% in all experiments and replicates). In previous studies, the decline in female fitness of ML selected haplotypes, combined with the masculinization of phenotype was treated as clinching evidence of IaSC. While this decline could also be due to a relaxation of selection acting upon the female-limited loci of the genome, we consider this possibility unlikely considering the magnitude and rapidity of decline and the consistency across replicate populations and experiments. More likely, such a consistent reduction in female fitness came about from sexually antagonistic costs. Fitness costs experienced by females carrying ML-selected haplotypes seems to be consistent across assay and evolved conditions, while male benefit variation might be specific to the local conditions of selection.

A few further observations underscore our concerns about specificity of the response to local conditions. Firstly, in the hemiclonal analysis of the ML_1_/MC_1_ population pair, we found a strong and significant difference in male fitness in the predicted direction (ML > MC by about 20%). Genetic variance for fitness was lower for ML_1_ compared to MC_1_, as predicted, with the data suggesting that selection had removed genotypes with low male fitness (fig. 4). However, an important caveat of these specific results is that the Y chromosomes were derived from the ML selection treatment (fig. S3), which were potentially altered by exposure to generations of maternal transmission. It is possible that the apparent improvement in fitness and reduction in genetic variance for male fitness in MLs actually results from inflated variance in the MCs that were exposed to these novel Y chromosomes. In the same experiment, mean female productivity and genetic variance for productivity was not different between ML_1_ and MC_1_. We note that for logistical ease we chose to assay female productivity here, rather than fitness under competitive conditions, which might explain the absence of a difference.

Overall, we find that more than 80 generations of male-limited selection resulted in dramatic improvements in male fitness under evolved conditions, but those increases are not consistently apparent when the evolved genomic haplotypes were tested in matched control conditions. This might reflect the exclusive recruitment of specialized compensatory and local adaptations, but this cannot explain the consistent reduction of fitness in females carrying ML hemiclonal genomes. Alternatively, specialized adaptations to the ML selection regime may have actually generated a cost under control (CRF) conditions, moving them off of their fitness peak with male fitness being locally “neutralized” and female fitness generally measureably reduced. In this case, sexually antagonistic variation was selected upon, but the fitness gains of males were cancelled out, leaving a signature only in reduced female fitness. We note that previous male limited selection experiments have not resolved potential contributions from the array of artefacts used in ML evolution: Rice (1996) (fitness tested under selection conditions), Rice (1998) (CG derived Y chromosomes, double dose haplotypes), Prasad *et al*. (2007) (translocated autosomes, CG derived Y chromosomes and CG females) and Abbott *et al*. (2013) (CG derived cytotypes, Y chromosomes and reversed sex chromosome inheritance).

We note the high concentration of research on *D. melanogaster* upon a single population (LH_m_), which may or may not be representative of the species as a whole. Our results also point to a need to further investigate the impact of specialized local and compensatory adaptation in the application of such selection protocols in order to understand their insights into sexual conflict. We echo Abbott et. al.’s (2023) lead off question: “Why is measuring and predicting fitness under genomic conflict so hard?”. In research currently *in prep*, we aim to answer many of the questions raised here by systematically varying all known artefacts involved in male- limited evolution.

## Author contributions

HT, TD and AKC developed the overall theme of the study; HT and AKC designed the experiments; HT, AKC, IS, MGB, & JAK carried out the experiments; HT analysed the data; HT wrote the first draft of the paper; HT, TD and AKC contributed to the final draft and revisions.

## Acknowledgments

Several undergraduate students were involved in lab maintenance and data collection during the period of this study. We thank Avery Want, Amanda Wigney, Navageevan Navamenon, Halle McNamara, Ronni Prince, Katelyn Viau, Alex Montenegro-Monreal, and Simmal Grewal for their efforts. We thank Karl Grieshop for his valuable inputs in interpreting these unexpected results. Funding for the work was from NSERC Discovery grants to AKC & TD.

## Data Accessibility Statement

Data archived here: osf.io/jgp59/

