## Supplement for "Mixed evidence for intralocus sexual conflict from male-limited selection in *Drosophila melanogaster*"

### Supplementary Material

Intralocus sexual conflict explored through male-limited selection in *Drosophila melanogaster*: A failure to replicate.

**Table S1.** The results of the ANOVA fit for the male CRF - generations 50, 64

| *Response* | | *Variable* | *df* | *F* | *P* |
|---|---|---|---|---|---|
| Fitness | Gen 50 | Selection | ***1*** | ***0.9711*** | ***0.33*** |
| | | Replicate | ***2*** | ***2.2390*** | ***0.12*** |
| | | Sel:Rep | ***2*** | ***1.2323*** | ***0.30*** |
| Fitness | Gen 64 | Selection | ***1*** | ***0.5599*** | ***0.46*** |
| | | Replicate | ***2*** | ***0.7995*** | ***0.46*** |
| | | Sel:Rep | ***2*** | ***0.7617*** | ***0.48*** |

**Table S2.** The results of the ANOVA fit for the male CRF - generation 70

| *Response* | | *Variable* | *df* | *F* | *P* |
|---|---|---|---|---|---|
| Fitness | Gen 70 | Treatment | ***1*** | ***7.3354*** | ***0.0003*** |
| | | Replicate | ***2*** | ***1.6979*** | ***0.19*** |
| | | Sel:Rep | ***2*** | ***2.2094*** | ***0.0543*** |

**Table S2a.** Contrast testing between levels of treatment in male CRF (gen 70) ANOVA. P value adjustment: Tukey method for comparing a family of 4 estimates

| *Contrast* | *Estimate* | *SE* | *t ratio* | *df* | *P* |
|---|---|---|---|---|---|
| MC:MLDD | ***0.0789*** | ***0.0244*** | ***3.237*** | ***68*** | ***0.0099*** |
| MC:MLSD | ***-0.0221*** | ***0.0244*** | ***-0.907*** | ***68*** | ***0.80*** |
| MC:MLSDa | ***0.0336*** | ***0.0244*** | ***1.378*** | ***68*** | ***0.52*** |
| MLDD:MLSD | ***-0.1010*** | ***0.0244*** | ***-4.144*** | ***68*** | ***0.0006*** |
| MLDD:MLSDa | ***-0.0453*** | ***0.0244*** | ***-1.858*** | ***68*** | ***0.26*** |
| MLSD:MLSDa | ***0.0557*** | ***0.0244*** | ***2.285*** | ***68*** | ***0.11*** |

**Table S3.** The results of the F test for male CRF heritable variance

| *Response* | *Variable* | | *MS* | *df* | *F* | *P* |
|---|---|---|---|---|---|---|
| Fitness | Line | MC | ***0.17705*** | ***37*** | ***3.4798*** | ***~0.0001*** |
| Fitness | Line | ML | ***0.05088*** | ***37*** | | |

**Table S4.** The results of the F test for female productivity heritable variance

| *Response* | *Variable* | | *MS* | *df* | *F* | *P* |
|---|---|---|---|---|---|---|
| Fitness | Line | ML | ***0.4670*** | ***37*** | ***1.1013*** | ***0.38*** |
| Fitness | Line | MC | ***0.4240*** | ***37*** | | |

**Table S5.** The results of ANOVAs on male CRF and female productivity modelled by selection.

| *Response* | *Variable* | *df* | *F* | *P* |
|---|---|---|---|---|
| Male CRF | Selection | ***1*** | ***17.766*** | ***<0.0001*** |
| Female productivity | Selection | ***1*** | ***1.1464*** | ***0.29*** |

**Table S6.** The results of intersex genetic correlation ($r_{mf}$) tests on male CRF and female productivity.

| | *Estimate* | *df* | *t* | *P* |
|---|---|---|---|---|
| ML | ***0.0247*** | ***36*** | ***0.1481*** | ***0.88*** |
| MC | ***0.1062*** | ***36*** | ***0.6408*** | ***0.53*** |

**Table S7.** The results of a two-tailed Fisher's Z test comparing $r_{mf}$ estimates from ML and MC lines

| | *Estimate* | z | *P* |
|---|---|---|---|
| MC-ML | ***0.0815*** | ***0.4195*** | ***0.34*** |

**Table S8.** The results of the ANOVA fit for the female CRF - generations 48, 50

| *Response* | | *Variable* | *df* | *F* | *P* |
|---|---|---|---|---|---|
| Fitness | Gen 48 | Selection | ***1*** | ***43.928*** | ***<0.0001*** |
| | | Replicate | ***2*** | ***2.0172*** | ***0.15*** |
| | | Sel:Rep | ***2*** | ***1.5259*** | ***0.23*** |
| Fitness | Gen 50 | Selection | ***1*** | ***91.803*** | ***<0.0001*** |
| | | Replicate | ***2*** | ***14.285*** | ***<0.0001*** |
| | | Sel:Rep | ***2*** | ***12.637*** | ***~0.0001*** |

**Table S8a.** Pairwise contrast testing between selection treatments within each level of replicate. Adjusted alpha rate for 3 comparisons is $\alpha = 0.0169$.

| *Response* | *Replicate* | *Variable* | *df* | *F* | *P* |
|---|---|---|---|---|---|
| Fitness | 1 | Selection | ***1*** | ***41.830*** | ***<0.0001*** |
| | 3 | Selection | ***1*** | ***68.547*** | ***<0.0001*** |
| | 5 | Selection | ***1*** | ***3.9771*** | ***0.0741*** |

**Table S9.** The results of the ANOVA fit on a generalized linear model (binomial error distribution) for male mating success.

| *Response* | *Variable* | *df* | *F* | *P* |
|---|---|---|---|---|
| Mating Success | Selection | ***1*** | ***2.0752*** | ***0.15*** |
| | Replicate | ***2*** | ***1.7469*** | ***0.18*** |
| | Sel:Rep | ***2*** | ***0.1189*** | ***0.88*** |

**Table S10.** The results of the ANOVA fit for male mating latency.

| *Response* | *Variable* | *df* | *F* | *P* |
|---|---|---|---|---|
| Mating Latency | Selection | ***1*** | ***1.4846*** | ***0.22*** |
| | Replicate | ***2*** | ***0.7299*** | ***0.48*** |
| | Sel:Rep | ***2*** | ***1.1045*** | ***0.34*** |

**Table S11.** The results of the ANOVA fit for male mating duration.

| *Response* | *Variable* | *df* | *F* | *P* |
|---|---|---|---|---|
| Mating Duration | Selection | ***1*** | ***0.2969*** | ***0.59*** |
| | Replicate | ***2*** | ***1.1592*** | ***0.32*** |
| | Sel:Rep | ***2*** | ***0.2941*** | ***0.75*** |

**Table S12.** The results of the ANOVA fit for fecundity induced by target males.

| *Response* | *Variable* | *df* | *F* | *P* |
|---|---|---|---|---|
| Fecundity induced | Selection | ***1*** | ***0.1000*** | ***0.75*** |
| | Replicate | ***2*** | ***1.0035*** | ***0.37*** |
| | Sel:Rep | ***2*** | ***0.3706*** | ***0.69*** |

**Table S13.** The results of the ANOVA fit for brood sex ratio of target male's offspring.

| *Response* | *Variable* | *df* | *F* | *P* |
|---|---|---|---|---|
| Sex ratio | Selection | ***1*** | ***0.0609*** | ***0.81*** |
| | Replicate | ***2*** | ***1.7600*** | ***0.18*** |
| | Sel:Rep | ***2*** | ***2.6786*** | ***0.0726*** |

**Table S14a.** The results of the ANOVA fit on a generalized linear model (binomial error distribution) on the number of target males that sired 100% of their mate's offspring.

| *Response* | *Variable* | *df* | *F* | *P* |
|---|---|---|---|---|
| WT.only Progeny | Treatment | ***1*** | ***1.5359*** | ***0.20*** |
| | Replicate | ***2*** | ***0.2647*** | ***0.77*** |
| | Sel:Rep | ***2*** | ***1.1248*** | ***0.35*** |

**Table S14b.** The results of the ANOVA fit for offspring sired, by target males that did not sire 100% of their mate's offspring.

| *Response* | *Variable* | *df* | *F* | *P* |
|---|---|---|---|---|
| P2 | Treatment | ***1*** | ***1.6598*** | ***0.20*** |
| | Replicate | ***2*** | ***2.2473*** | ***0.11*** |
| | Sel:Rep | ***2*** | ***1.3440*** | ***0.26*** |

**Figure S1.** (a) ML selection breeding design. (b) Recombination box design. CG females are denoted as DTW or DTP based on presence or absence of *bw*$^D$ marker.

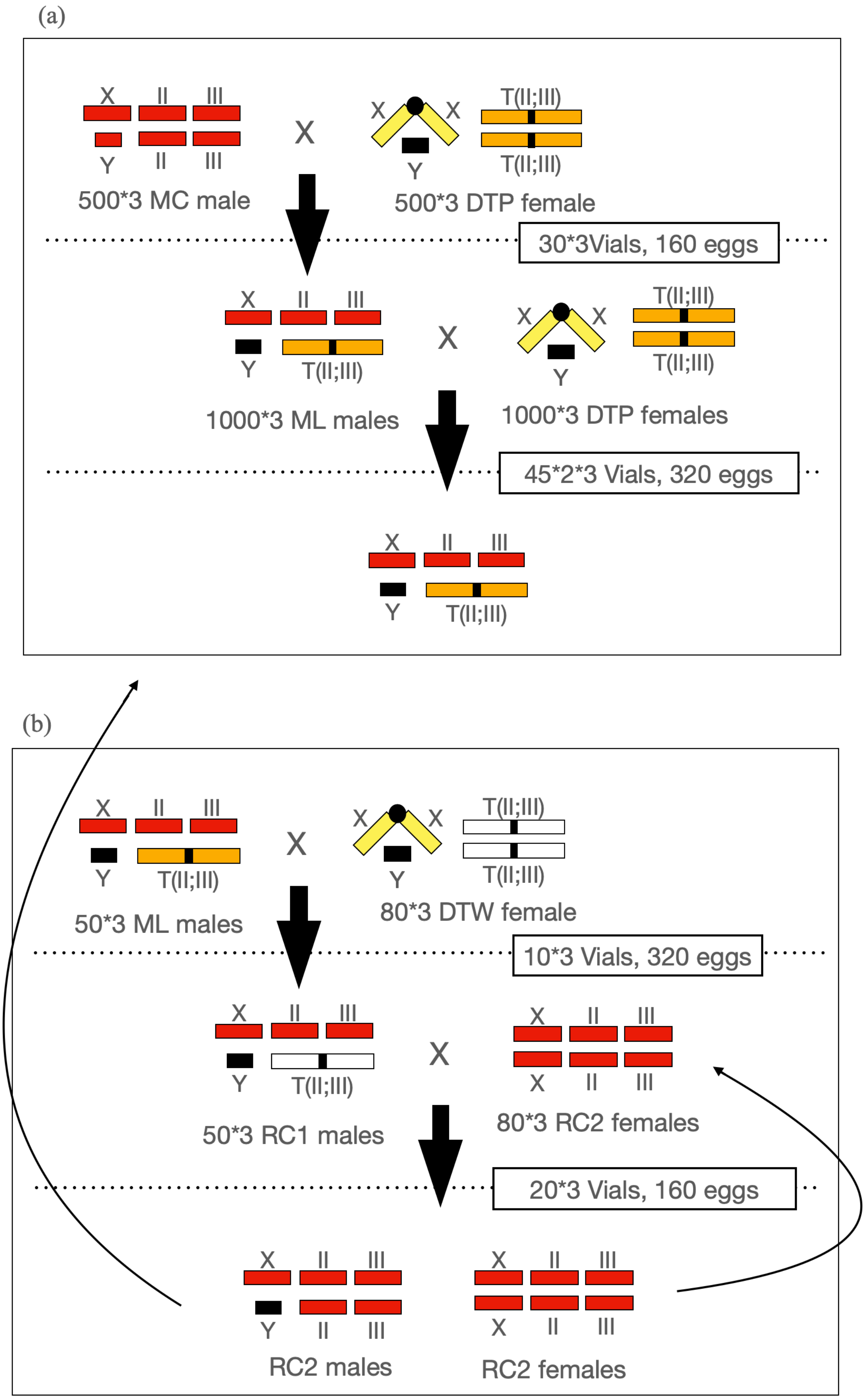

**Figure S2.** CRF experimental males:

(a)

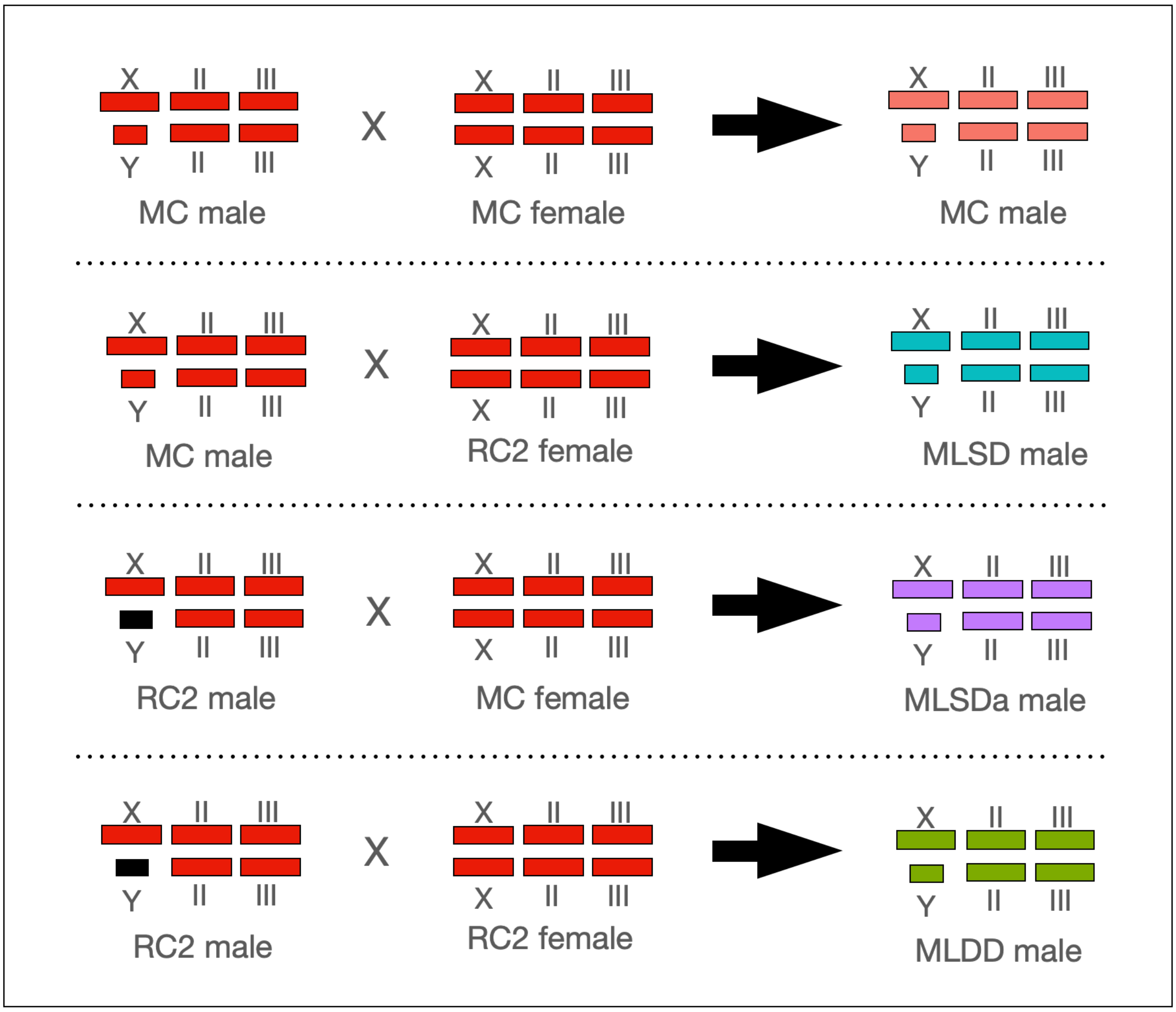


(b)

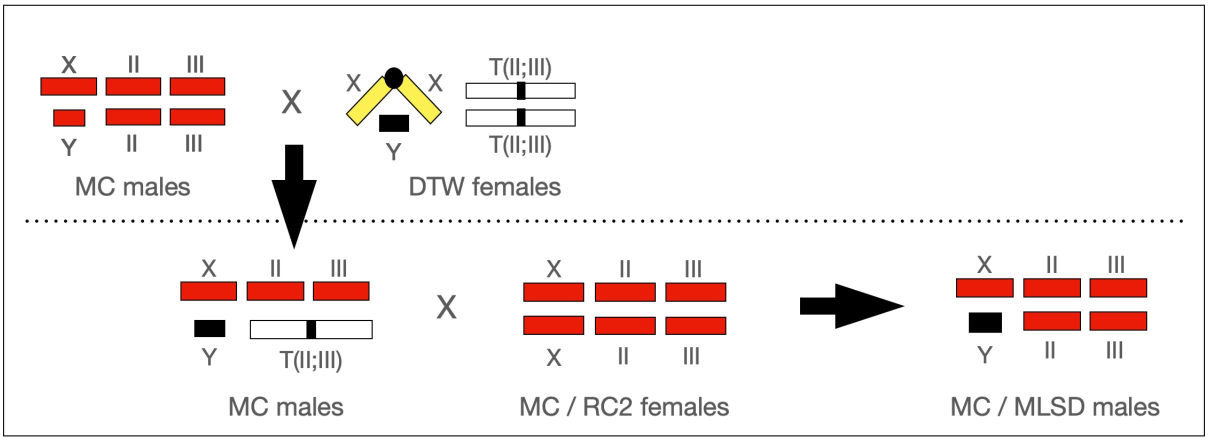

**Figure S3.** Hemiclonal analysis – breeding design

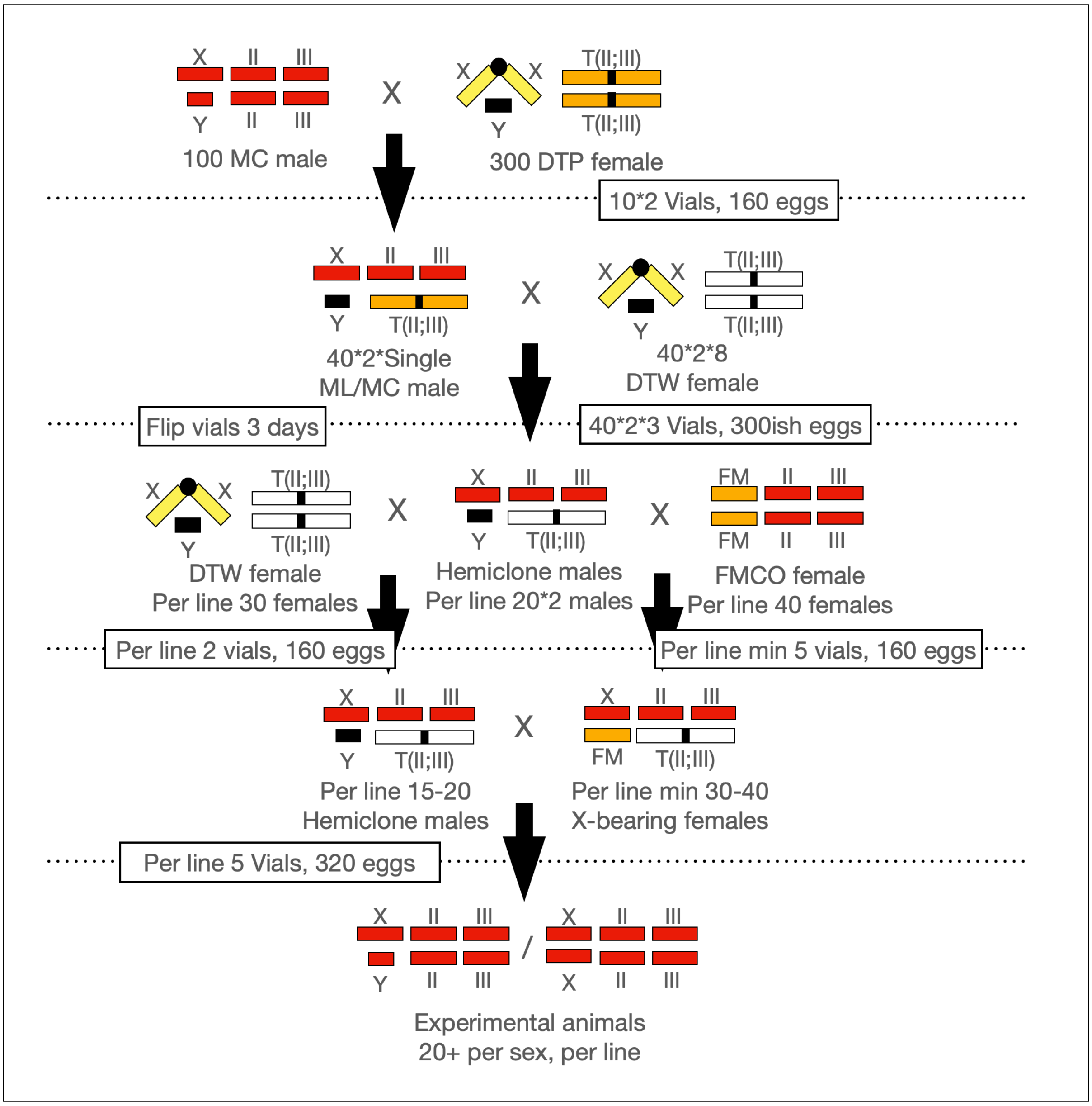
